# Evaluation of the rapidBACpro^®^ II kit for the rapid identification of microorganisms directly from blood cultures using MALDI-TOF MS

**DOI:** 10.1101/2021.01.25.428200

**Authors:** Marina Oviaño, André Ingebretsen, Anne K Steffensen, Antony Croxatto, Guy Prod’hom, Lidia Quiroga, Germán Bou, Gilbert Greub, Belén Rodríguez-Sánchez

## Abstract

**Objectives:** Identification of microorganisms directly from blood cultures (BCs) using MALDI-TOF MS has shown to be the application with most impact in this methodology. In this study, a novel commercial method, the rapidBACpro^®^ II, was evaluated in four clinical microbiology laboratories.

**Methods:** Positive blood culture samples (n=801) were processed using the rapidBACpro^®^ II kit and then compared with routine gold standard. A subset of monomicrobial BCs (n=560) were analyzed in parallel with the Sepsityper^®^ kit (Bruker Daltonics, Bremen, Germany) and compared with the rapidBACpro^®^ II kit. In addition, the rapidBACpro^®^ II kit was also compared with two different in-house methods.

**Results:** Overall, 80.0% of the monomicrobial isolates (609/761) were correctly identified by the rapidBACpro^®^ II kit at the species level (92.3% of the Gram negative and 72.4% of the Gram positive bacteria). The comparison with the Sepsityper^®^ kit yielded higher rates of correct species-level identification provided by the rapidBACpro^®^ II kit for all categories (p>0.0001) except for yeasts identified with score values >1.7. It also proved superior to the ammonium chloride method (p>0.0001) but the differential centrifugation method allowed higher rates of correct identification for Gram negative bacteria (p>0.1).

**Conclusions:** The rapidBACpro^®^ II kit allowed a high rate of microorganisms correctly identified. The percentage of accurate species-level identification of Gram positive bacteria was particularly noteworthy in comparison with other commercial and in-house methods. This fact was especially interesting in the case of *Staphylococcus* sp. and *Streptococcus* sp. in order to elucidate their clinical impact, for example in device-associated bacteremia.

## INTRODUCTION

Rapid identification of microorganisms causing bloodstream infections (BSI) is one of the applications with most impact of MALDI-TOF Mass Spectrometry (MS) in clinical microbiology (1, 2). This technology has demonstrated timely and accurate identification of a wide variety of microorganisms directly from positive blood cultures (BCs) and, in addition, it is inexpensive and efficient (3). Most microbiology laboratories worldwide have implemented this approach in order to guide clinicians to initiate early optimal antimicrobial treatment (4), a factor that has been correlated with higher rates of positive outcomes (5).

Different pre-processing methods have been reported for the successful identification of microorganisms directly from positive blood cultures using MALDI-TOF MS. Most of them focus on the separation of the microorganisms present in blood cultures either using a differential centrifugation method that eliminates blood cells and other contaminants (6) or by an improved erythrocyte-lysing procedure that employs ammonium chloride (7), sulfate dodecyl sodium (8) or saponin (9). Other studies have reported short-incubations of BC broth on agar plates (2 to 6 hours) prior to MALDI-TOF MS identification from the velum grown on the surface of the plate (10). This proceeding has allowed successful identification of bacteria and even helped overcome some of the limitations of MALDI-TOF MS: accurate identification of more than one pathogen in polymicrobial infections and reliable identification of gram positive microorganisms (11).

Commercial kits have been manufactured for the improved lysis of blood cells and subsequent recovery of the microorganisms present in positive BCs, facilitating the standardization and implementation of this task in the routine of the clinical microbiology laboratory. Most studies have reported the successful identification of microorganisms using the Sepsityper^®^ kit (Bruker Daltonics, Bremen, Germany) (12) and, more recently, the Vitek^®^ MS Blood Culture kit (bioMérieux, Lyon, France) (13). However, new chemistries have been developed. In this study, a new available method containing a polyallylamine-polystyrene copolymer, the rapidBAC pro^®^ II kit (Nittobo Medical Co., Tokyo, Japan), has been evaluated in four research centers. The performance of this commercial kit was compared with the Sepsityper^®^ kit and additionally with two in-house methods: differential centrifugation and lysis of blood cells with ammonium chloride (Figure 1).

**Figure 1.**
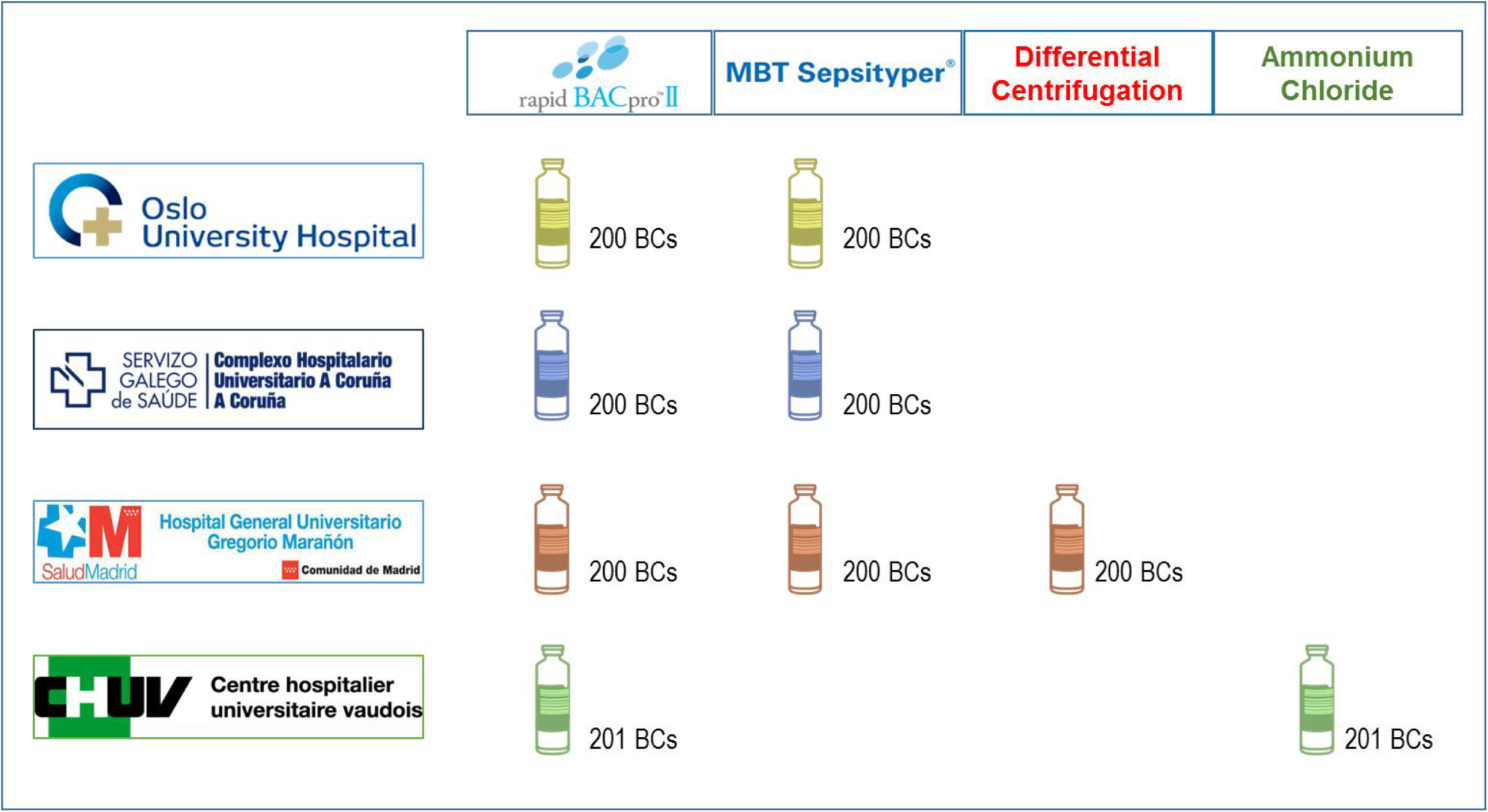
Study design. 200 BCs were collected in each laboratory -201 in the CHUV- and analyzed using the rapidBAC pro^®^ II kit. The results were compared with the Sepsityper^®^ kit (CHUAC, HGM and OUH), with the ammonium-chloride lysis system (CHUV) and with the differential centrifugation method (HGM).

## MATERIALS AND METHODS

### Study settings

The evaluation of the rapidBACpro^®^ II kit (Nittobo Ltd., Tokyo, Japan) has been carried out in four research centers: the Microbiology Department from Complejo Hospitalario Universitario A Coruñ a –CHUAC- (A Coruñ a, Spain), the Department of Microbiology at Oslo University Hospital –OUH-(Oslo, Norway)-, the Institute of Microbiology from the University Hospital of Lausanne–CHUV- (Lausanne, Switzerland), and the Clinical Microbiology and Infectious Diseases Department from the Gregorio Marañón University Hospital –HGM- (Madrid, Spain) and. In each laboratory, 200 consecutive positive blood cultures –belonging to 200 (CHUAC), 200 (OUH), 190 (HGM) and 201 (CHUV) patients-were collected between January and June 2019. Aerobic and anaerobic blood culture bottles (BD BACTEC Plus Aerobic/F and Plus Anaerobic/F Culture Vials, Becton Dickinson, Franklin Lakes, NJ, USA) were incubated at 35°C for up to 5 days in the BACTEC™ FX incubation system (Becton Dickinson). When the bottles flagged positive, Gram staining was performed and a small amount of broth was cultured on suitable agar media and further incubated at 35°C overnight. Growth on agar media and the subsequent identification by MALDI-TOF MS was considered as a gold standard. In parallel, the evaluation of the rapidBACpro^®^ II kit was carried out.

The performance of the rapidBACpro^®^ II kit (Nittobo Ltd.) was compared with that of the Sepsityper^®^ kit (Bruker Daltonics, Bremen, Germany) at 3 centers (CHUAC, HGM and OUH, n=600) At one center (CHUV) the rapidBACpro^®^ II kit was compared with an in-house processing method (7) which employs ammonium chloride (n=201) In addition, the HGM laboratory also compared the rapidBACpro^®^ II kit with their in-house differential centrifugation method(n=200) (2, 6) (Figure 1).

### Sample preparation using the rapidBACpro^®^ II kit

Bacterial pellets from positive blood cultures were processed following the manufacturer’s instructions. Briefly, 1ml of broth was mixed with 500μl of lysis buffer and the mixture was centrifuged for 3 minutes at 2000-5000g. The supernatant was thoroughly removed; the pellet was resuspended in 800μl of deionized water and transferred to another tube containing 200μl of cationic polymer and 200μl of reaction buffer, previously mixed. The mixture was vortexed and centrifuged at 2000-5000g for 1 minute. Then, the supernatant was removed again and the pellet resuspended in 800μl of 70% ethanol. After complete resuspension of the pellet by pipetting it up and down, the sample was centrifuged again in the above-mentioned conditions and the pellet was submitted to a protein extraction step with 30μl of formic acid 70% and the same amount of acetonitrile. After another centrifugation step, 1-2μl of supernatant were deposited on the MALDI target plate, allowed to dry and covered with 1μl of HCCA matrix, prepared according to the manufacturer instructions (Bruker Daltonics, Bremen, Germany)

### Sample preparation using the Sepsityper® kit

Sample preparation of bacterial pellets was performed according to the instructions of the manufacturer (Bruker Daltonics, Bremen, Germany). The method consisted of mixing 1ml of blood culture broth with 200μl of lysis buffer. The mix was centrifuged for 2 min at 13000 rpm. The pellet was washed with 1ml of washing buffer and centrifuged for 1 min at 13000 rpm. The supernatant was removed and the pellet submitted to protein extraction with formic acid and acetonitrile. As explained above, 1-2μl of supernatant were transferred onto the MALDI target plate and covered with 1μl of HCCA matrix.

### Ammonium chloride erythrocyte-lysing procedure

Ammonium chloride erythrocyte-lysing procedure was applied as previously described (7). Five ml of positive blood-culture broth was resuspended in 45 ml of deionized water and centrifuged at 1000 g for 10 minutes at room temperature. The supernatant containing lysed blood-cells was discarded and the pellet was suspended in 1 ml of ammonium chloride (0.15 M NH4Cl, 1 mM KHCO3, pH 7.31). A second centrifugation step at 140 g for 10 min was applied and the supernatant discarded. The final pellet was suspended in 0.2 ml of deionized water and 1-2 μl was transferred to the MALDI target plate and, once dry, covered with formic acid and, subsequently with HCCA matrix.

### Differential centrifugation method

From each BC, 8 ml of broth were transferred to a 15 ml tube and centrifuged at 150 g for 10 min. The supernatant was then collected in four 1.5-ml tubes and centrifuged again at 13000 rpm for 1 min. The pellets from the four tubes were collected in one tube and washed with 1ml deionized water. After another centrifugation step under the same conditions, the supernatant was discarded and the pellet spotted directly onto the MALDI target plate. The spots were allowed to dry at room temperature and then covered with 1μl of formic acid. Once dried, HCCA matrix was added (2).

### Identification of by MALDI-TOF MS

MALDI-TOF MS analysis was performed on a Bruker Microflex LT mass spectrometer using the MBT Compass Library (#1829023) containing 7331 reference spectra. FlexControl 3.3 and Maldi Biotyper 3.0 software (Bruker Daltonics) were applied to acquire the spectra and for the identification of the isolates with the standard Biotyper module, respectively. Bacterial Test Standard (Bruker Daltonics) was used for calibration purposes. The identification was performed in duplicates and the highest score from each pair was recorded.

### Interpretation of the results

In this study, score values ≥2.0 and ≥1.7 were established as the cut-off value for species- and genus-level identification, respectively. Isolates identified with score values below 1.6 were taken into account only when the first three identifications provided by MALDI-TOF MS were consistent. Otherwise, the identification was considered “not reliable”.

### Statistical analysis

Sensitivity values for each pre-processing method were calculated. They were compared with the sensitivity of the rapidBACpro^®^ II kit using the McNemar test for paired samples with two tails. Validity values were calculated with a 95% confidence interval (CI) following an exact binomial distribution.

## RESULTS

### Performance of the rapidBACpro^®^ II kit

During the study period, 801 consecutive, positive BCs were collected in the 4 participating laboratories and the present microorganisms were identified by MALDI-TOF MS after sample pre-processing using the rapidBAC pro^®^ II kit. In 761 BCs -560 from CHUAC, OUH and HGM and 201 from CHUV-, only one species was present (Table 1) while in 27 BCs more than one microorganism was present (Supplementary Table 1). In the remaining BCs (n=13) no growth were recorded after 24 h incubation on agar plates, despite flagging positive in their BACTEC systems. The Gram staining of these BCs were reviewed and considered as inconclusive in 13/13 cases. Therefore, these BCs were considered as false positives and removed from the study.

**Table 1.** Identification by MALDI-TOF MS using the rapidBACpro^®^ II kit of all the monomicrobial BCs analyzed in the four participating laboratories.

|  |  | n | rapidBACpro II test |  |  |  |  |  |
| --- | --- | --- | --- | --- | --- | --- | --- | --- |
|  |  |  | S >2.0 | (%) | 1.7 < S < 1.99 | (%) | S < 1.7 | (%) |
| (A) |  |  |  |  |  |  |  |  |
|  | Overall | 761 | 609 | 80.0 | 110 | 14.4 | 42 | 5.6 |
|  | Gram-Negative (GN) | 323 | 298 | 92.3 | 20 | 6.2 | 5 | 1.5 |
|  | Gram-Positive (GP) | 417 | 302 | 72.4 | 86 | 20.6 | 29 | 7.0 |
| (B) |  |  |  |  |  |  |  |  |
|  | Enterobacteriales | 271 | 254 | 93.7 | 15 | 5.5 | 2 | 0.8 |
|  | *Escherichia coli* | 149 | 139 | 93.3 | 9 | 6.0 | 1 | 0.7 |
|  | *Klebsiella* spp | 70 | 68 | 97.1 | 2 | 2.9 | 0 | 0 |
|  | *Pseudomonas* | 15 | 15 | 100 | 0 | 0 | 0 | 0 |
|  | Other GN | 37 | 29 | 78.4 | 5 | 13.5 | 3 | 8.1 |
|  | *S. aureus* | 95 | 90 | 94.7 | 2 | 2.1 | 3 | 3.2 |
|  | CoN staphylococci | 153 | 91 | 59.5 | 53 | 34.6 | 9 | 5.9 |
|  | Streptococci | 86 | 57 | 66.3 | 16 | 18.6 | 13 | 15.1 |
|  | Enterococci | 37 | 36 | 97.3 | 1 | 2.7 | 0 | 0 |
|  | Other GP | 46 | 28 | 60.9 | 14 | 30.4 | 4 | 8.7 |
|  | Yeast | 21 | 9 | 42.9 | 4 | 19.0 | 8 | 38.1 |

The implementation of the rapidBAC pro^®^ II kit in the four participating laboratories allowed an overall rate of 80.0% (609/761) correct identification at the species level (score values ≥2.0) of the pathogens present in the monomicrobial BCs (Table 1). Among different bacterial groups, this rate ranged between 66.7% and 89.0% (Supplementary Table 1). The identification at the genus level (scores ranging between 2.0 and 1.7) was accurate in another 110/761 isolates (14.4%). In only 42 cases the identification score values were below 1.7 (5.6%; 2.0-9.0% inter-laboratory range). This group of microorganisms was mainly composed of Gram positives (29/42) and yeasts (8/42), confirming the limitations of MALDI-TOF MS identification when applied directly to blood culture broth. Despite the low-confidence scores, these identifications were in agreement with the gold standard method (Table 1). Of the Gram negatives 92.3% were identified with score values ≥2.0 (298/323; 81.5-96.4% inter-laboratory range). The number of accurate identification at the species level is above 80.0% for all the main groups of bacteria within this category.

### Comparison between the rapidBAC pro^®^ II kit and the Sepsityper^®^ kit

A subset of 560 monomicrobial BCs were analyzed in 3 laboratories (CHUAC, HGM and OUH –Figure 1-) with both the rapidBAC pro^®^ II kit and the Sepsityper^®^ kit in a head-to-head evaluation (Table 2). Overall, 76.8% (n=430) of the BCs were identified at the species-level with the rapidBAC pro^®^ II kit vs. 69.3% (n=388) with the Sepsityper^®^ kit. Similar amount of BCs was identified at the genus level using both kits (16.4% -n=92-rapidBAC pro^®^ II kit vs. 15.2% -n=85-Sepsityper^®^ kit), whilst the rate of unreliable identifications was higher with the Sepsityper^®^ kit (6.8% -n=38- vs. 15.5% -n=87-) -Figure 2-. Gram negatives were successfully identified at the species level with both kits (90.5% (n=210) rapidBAC pro^®^ II kit vs. 85.8% (n=199) Sepsityper^®^ kit). Interestingly, differences were observed in the number of unreliable identifications (1.7% -n=4- rapidBAC pro^®^ II kit vs. 10.8% -n=25- Sepsityper^®^ kit). Regarding Gram positives, the rate of species-level identifications yielded by the rapidBAC pro^®^ II kit was higher (68.7% -n=215- vs. 59.7% -n=187-) than the Sepsityper^®^ kit and the number of BCs with score values below 1.7 was reduced (8.3% -n=26- vs. 17.9% -n=56-). The proportion of yeasts identified at the genus level, however, was higher using the Sepsityper^®^ kit (46.7% -7/15- vs. 13.3% -2/ 15-) and the number of unreliable isolates was lower (40.0% -n=6- vs. 53.4% -n=8-) although species-level identification was achieved more often with the rapidBAC pro^®^ II kit (33.3% -n=5- vs. 13.3% -n=2-) -Table 2-. Overall, the rapidBAC pro^®^ II kit showed higher sensitivity both at the species- (p>0.0001, OR=0.429 (0.287-0.630), CI 95%) and the genus-level (p>0.0001, OR=0.227 (0.125-0.388), CI 95%).

**Table 2.** Head-to-head comparison of the performance of both commercial kits on a set of monomycrobial BCs (n=560) analyzed in three laboratories (CHUAC, HGM and OUH).

|  |  | n | rapidBACpro II test |  |  |  |  |  | Sepsityper kit |  |  |  |  |  |
| --- | --- | --- | --- | --- | --- | --- | --- | --- | --- | --- | --- | --- | --- | --- |
|  |  |  | S >2.0 | (%) | 1.7 < S < 1.99 | (%) | S < 1.7 | (%) | S >2.0 | (%) | 1.7 < S < 1.99 | (%) | S < 1.7 | (%) |
| (A) |  |  |  |  |  |  |  |  |  |  |  |  |  |  |
|  | Overall | 560 | 430 | 76.8 | 92 | 16.4 | 38 | 6.8 | 388 | 69.3 | 85 | 15.2 | 87 | 15.5 |
|  | Gram-Negative (GN) | 232 | 210 | 90.5 | 18 | 7.8 | 4 | 1.7 | 199 | 85.8 | 8 | 3.4 | 25 | 10.8 |
|  | Gram-Positive (GP) | 313 | 215 | 68.7 | 72 | 23.0 | 26 | 8.3 | 187 | 59.7 | 70 | 22.4 | 56 | 17.9 |
| (B) |  |  |  |  |  |  |  |  |  |  |  |  |  |  |
|  | Enterobacteriales | 194 | 179 | 92.3 | 14 | 7.2 | 1 | 0.5 | 179 | 92.3 | 5 | 2.6 | 10 | 5.1 |
|  | Escherichia coli | 112 | 103 | 92.0 | 8 | 7.1 | 1 | 0.9 | 105 | 93.7 | 2 | 1.8 | 5 | 4.5 |
|  | Klebsiella spp | 43 | 41 | 95.3 | 2 | 4.7 | 0 | 0 | 41 | 95.4 | 1 | 2.3 | 1 | 2.3 |
|  | Pseudomonas | 10 | 10 | 100 | 0 | 0 | 0 | 0 | 7 | 70.0 | 0 | 0 | 3 | 30.0 |
|  | Other GN | 28 | 21 | 75.0 | 4 | 14.3 | 3 | 10.7 | 13 | 46.4 | 3 | 10.7 | 12 | 42.9 |
|  | S. aureus | 75 | 70 | 93.3 | 2 | 2.7 | 3 | 4.0 | 68 | 90.7 | 2 | 2.7 | 5 | 6.6 |
|  | CoN staphylococci | 118 | 66 | 55.9 | 44 | 37.3 | 8 | 6.8 | 51 | 43.2 | 36 | 30.5 | 31 | 26.3 |
|  | Streptococci | 67 | 39 | 58.2 | 15 | 22.4 | 13 | 19.4 | 34 | 50.7 | 21 | 31.3 | 12 | 18.0 |
|  | Enterococci | 30 | 29 | 96.7 | 1 | 3.3 | 0 | 0 | 23 | 76.7 | 4 | 13.3 | 3 | 10.0 |
|  | Other GP | 23 | 11 | 47.8 | 10 | 43.5 | 2 | 8.7 | 11 | 47.8 | 7 | 30.4 | 5 | 21.8 |
|  | Yeast | 15 | 5 | 33.3 | 2 | 13.3 | 8 | 53.4 | 2 | 13.3 | 7 | 46.7 | 6 | 40.0 |

**Figure 2.**
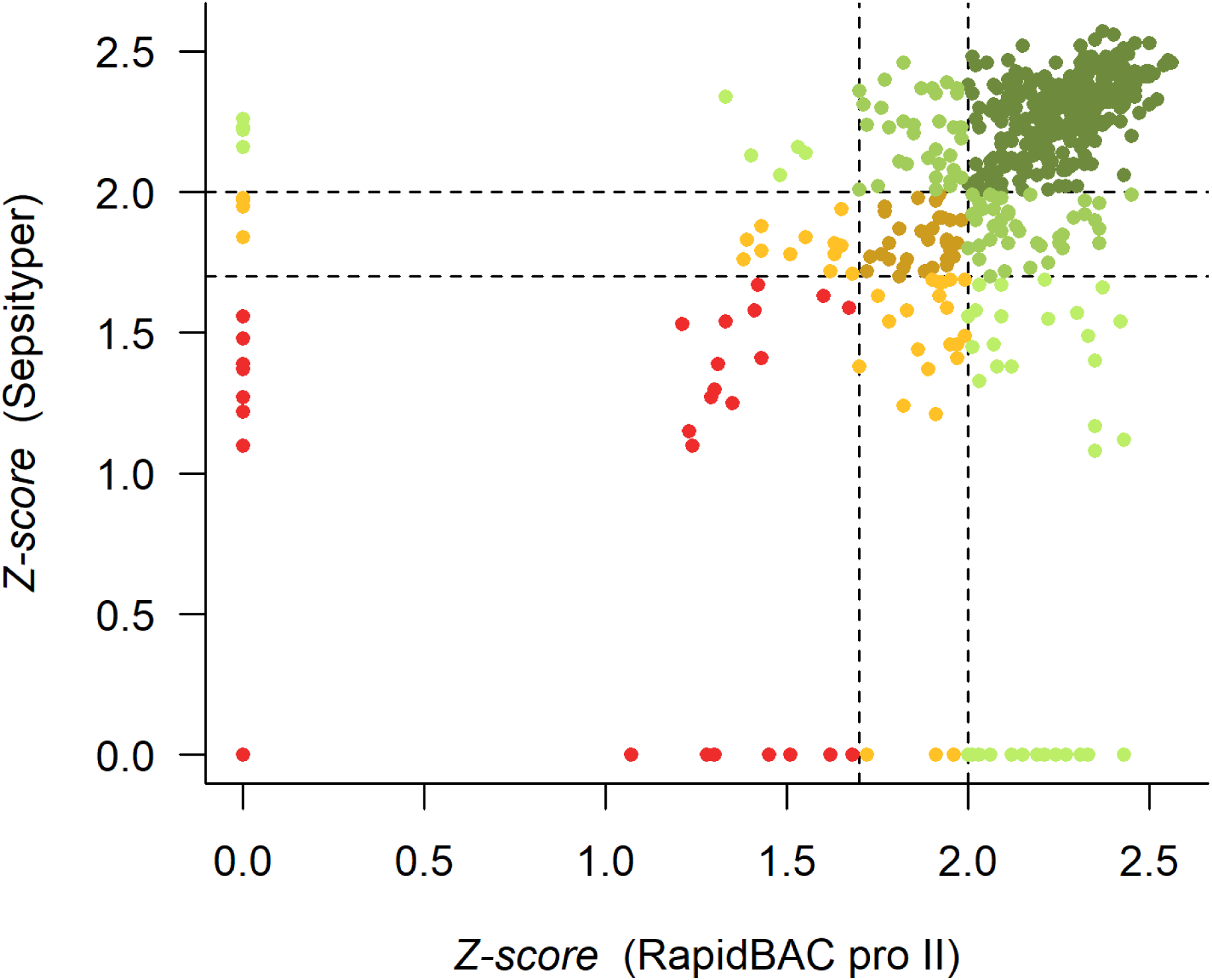
Scattering plot representing the distribution of the scores obtained by a group of 560 monomicrobial BCs analyzed using the rapidBAC pro^®^ II (in the x-axis) and the Sepsityper^®^ kit (in the y-axis). Dark green represents isolates identified at the species level by both methods; in green are represented those identified at the species level by one method and at the genus level by the other; isolates identified at the species level by one method and unreliably by the second method (score value < 1.7) are shown in light green. Similarly, isolates identified at the genus level by both methods are shown in brown; in yellow are represented those identified at the genus level by one method and unreliably by the second method. Isolates unreliably identified by both methods are shown in red.

In this study we also included 27 polymicrobial BCs. (Supplementary Table 2). Both commercial kits allowed correct species-level assignment of one microorganism in 23/27 BCs with score values ranging between 1.67 and 2.55 for the rapidBAC pro^®^ II kit and in 22/27 cases with the Sepsityper^®^ kit, with scores between 1.75 and 2.45. In two cases, both pathogens present in the same BC could be identified using the rapidBAC pro^®^ II kit (*Staphylococcus epidermidis* plus *Staphylococcus capitis* and *Klebsiella pneumoniae* plus *Staphylococcus haemolyticus*) and in a different case with the Sepsityper^®^ kit (*Staphylococcus aureus* plus *Enterococcus faecium*). Finally, both kits failed to identify all microorganisms from the same BC in two cases (*Capnocytophaga sputigena* plus *Fusobacterium nucleatum* and *S. aureus* plus *Streptococcus mitis*) and the Sepsityper^®^ kit also failed in two more cases where two microorganisms were present (Supplementary Table 1).

### Comparison between the rapidBAC pro^®^ II kit and the ammonium chloride method

The CHUV laboratory also compared the performance of rapidBAC pro^®^ II kit and their in-house method that employs ammonium chloride for cell lysis (7) in a head-to-head assay (Table 3). This analysis showed that the commercial kit allowed correct identifications at the species level with higher accuracy than the in-house method (89.0% -179/201- vs. 39.8% -80/201-; p>0.01) –Figure 3-. The rate of correct species-level assignment of the rapidBAC pro^®^ II kit was higher for both Gram negatives (96.7% -88/91- vs. 53.8% -49/91-) and Gram positives (83.6% -87/104- vs. 26.9% -28/104-). In the first case, the application of the commercial kit allowed the identification of 97.4% of the microorganisms from the *Enterobacteriaceae* family (75/77) and 100% of the *Pseudomonas* (5/5) vs. 61.0% (47/77) and 0% (0/5), respectively, when the in-house method was applied. In the case of the Gram positives, the ammonium chloride method allowed 40.0% species-level identification of *Staphylococcus aureus* (8/20) vs. 100% using the rapidBAC pro^®^ II kit (Table 3). The same is true for other groups of Gram positive microorganisms (Coagulase negative –CoN-Staphylococci, Staphylococci and Enterococci) for which the rapidBAC pro^®^ II kit yielded 71.4-100% accurate species-level identification, whilst the ammonium chloride method ranged between 26.4-28.6% correct identification at the same level. Similarly, the ammonium chloride method yielded high rates of unreliable identifications for both groups of microorganisms: 19.8% of the Gram negatives (18/91) and 38.5% of the Gram positives (40/104) were identified by MALDI-TOF MS with score values ≤1.7 whilst the use of the rapidBAC pro^®^ II kit reduced the amount of non-reliable identifications to 1.1% of the Gram negatives (1/91) and 2.9% of the Gram positives (3/104) -Table 3-. Likewise, the identification of yeasts at the species level was accurate at the species level for a higher number of isolates when the commercial kit was applied: 66.7% (4/6) vs. 50.0% (3/6) with the in-house method.

**Table 3.**
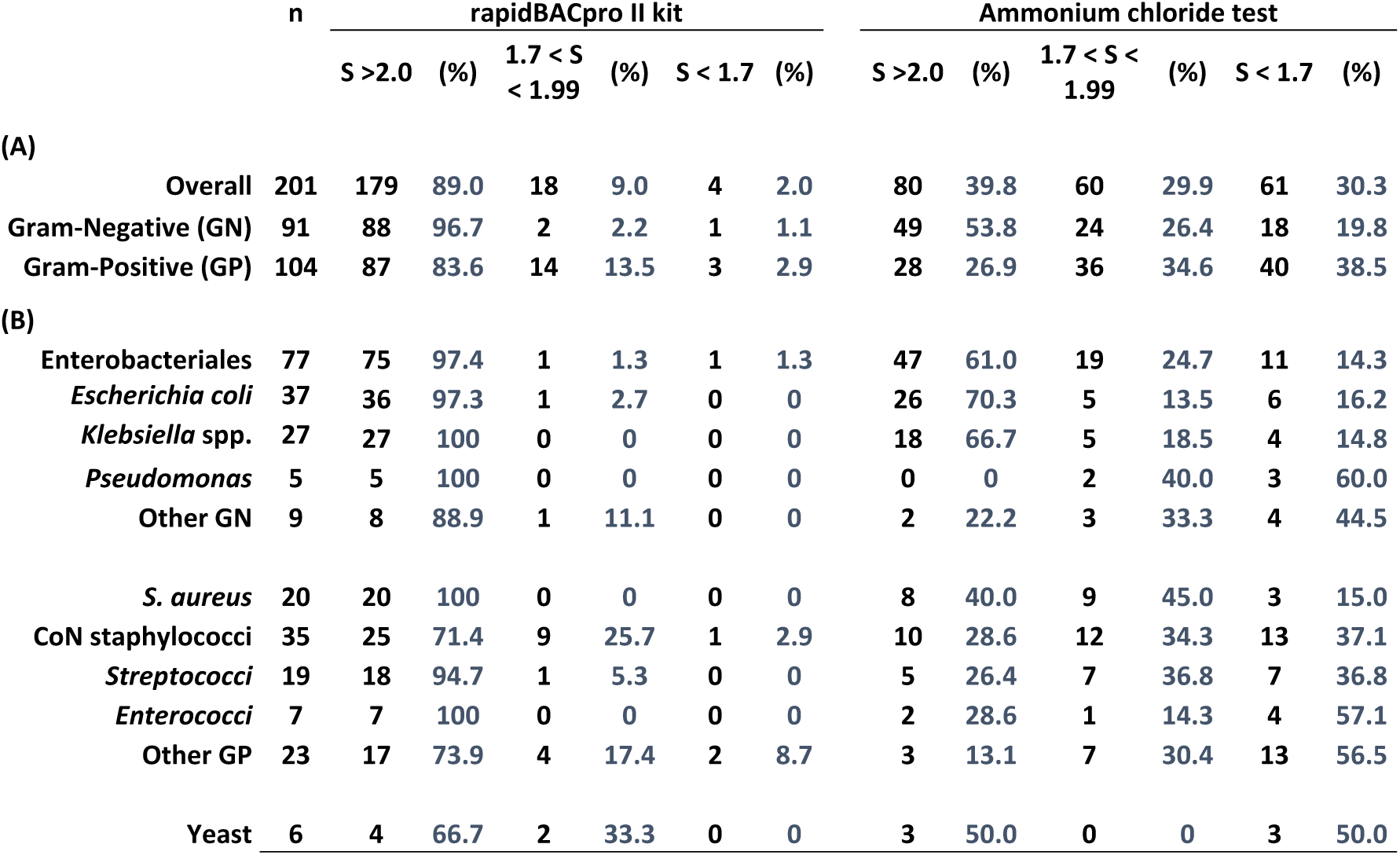
Head-to-head comparison between the performances of the rapidBACpro^®^ kit and the in-house method described by Prod’hom *et al*. (2010) that uses ammonium chloride.

**Figure 3.**
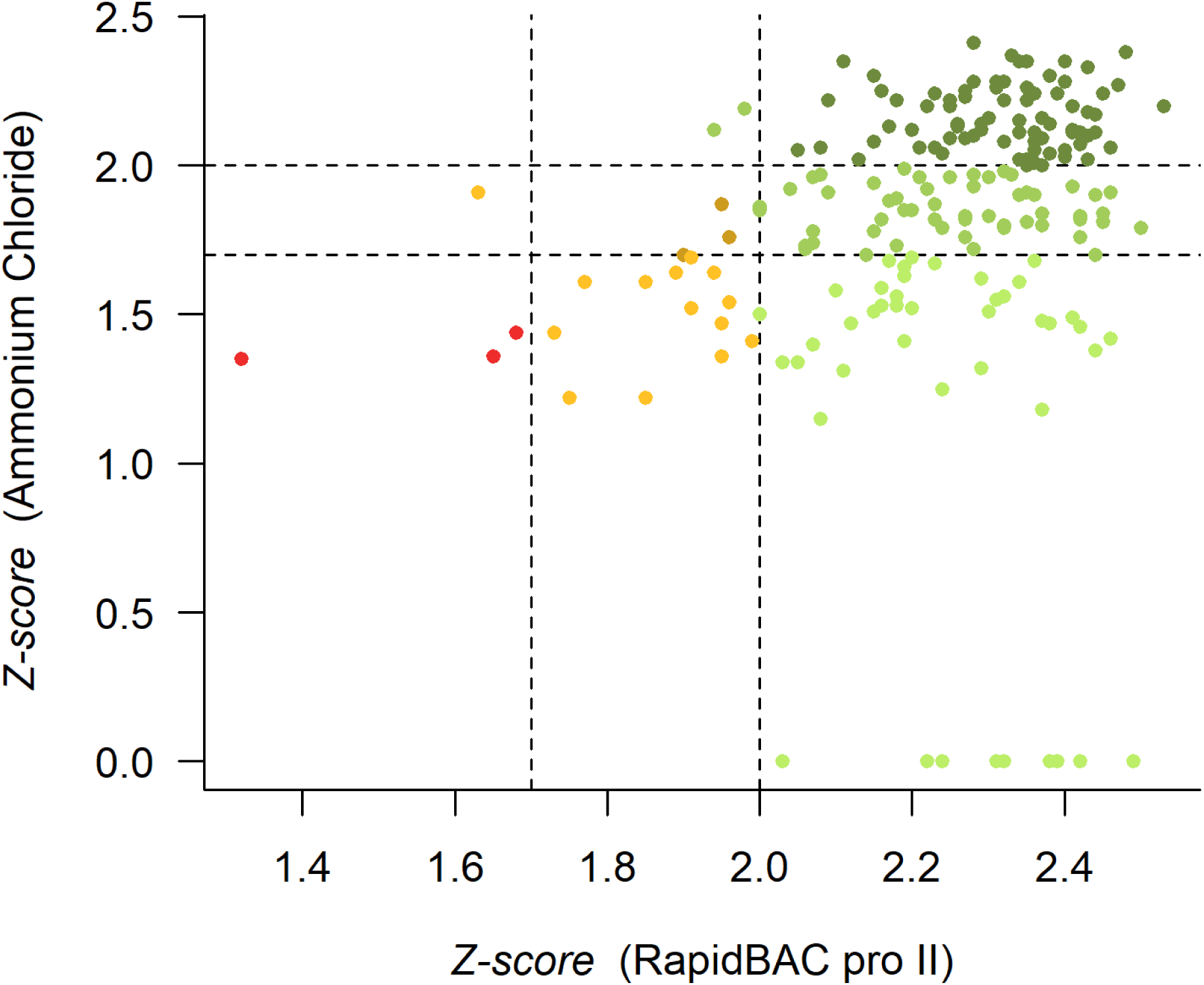
Representation of the scores from 201 BCs analyzed by applying the the rapidBAC pro^®^ II (in the x-axis) and the ammonium chloride method (in the y-axis). The same color code as in Figure 2 is applied.

**Figure 4.**
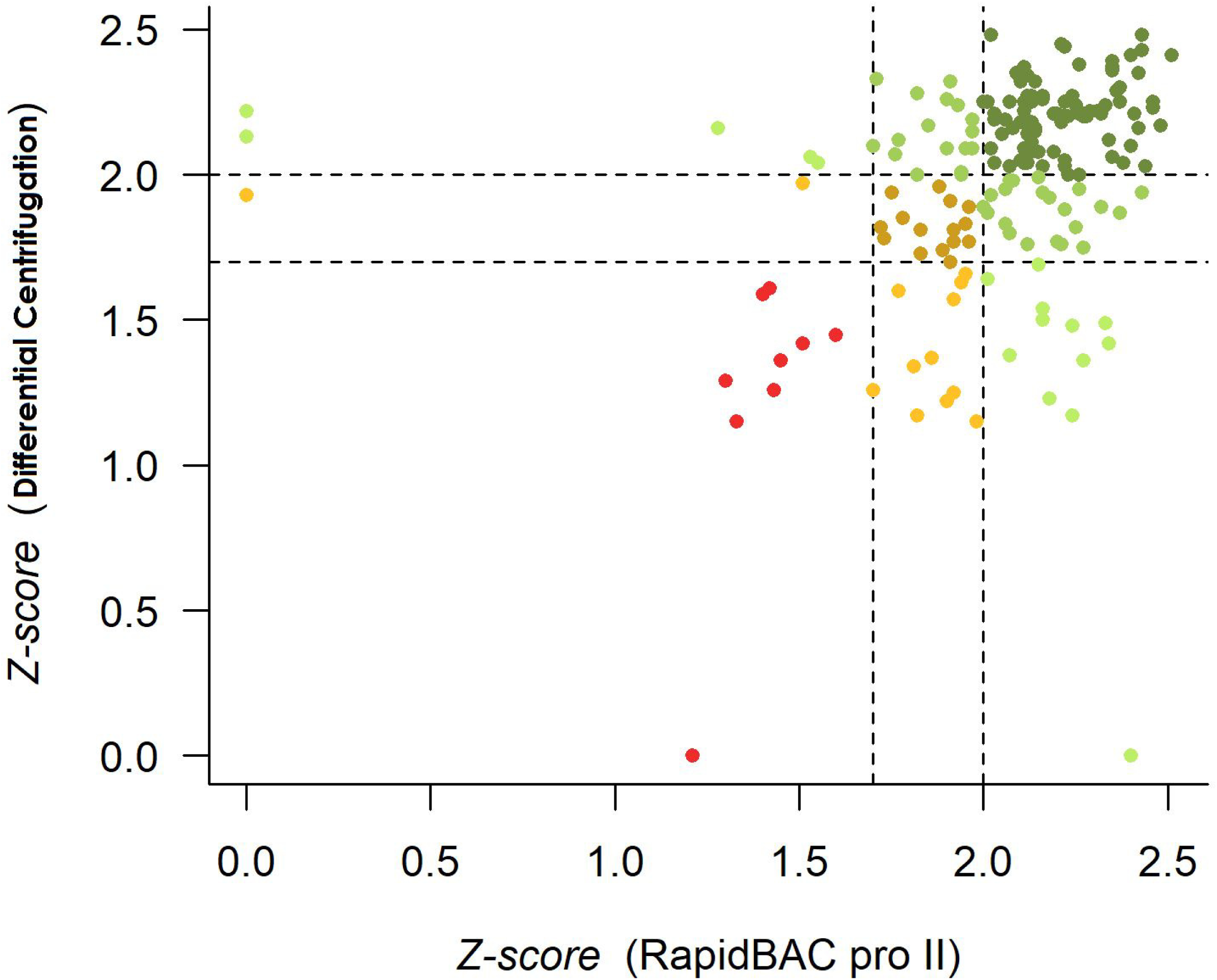
Scattering plot showing the distribution of the scores from 177 monomicrobial BCs analyzed using the rapidBAC pro^®^ II (in the x-axis) and the differential centrifugation method (in the y-axis). The same color code as in Figure 2 is applied.

### Comparison between the rapidBAC pro^®^ II kit and the differential centrifugation method

The performance of the rapidBAC pro^®^ II kit was also compared with the differential centrifugation in the HGM laboratory (Table 4). Overall, the commercial method allowed 66.7% (118/177) correct species-level assignment vs. 61.6% (109/177) obtained by using the in-house kit (p=0.14; OR=1.54 -0.88-2.77- CI 95%) –Figure 3-. Differences in the identification of Gram negatives and Gram positives were observed. The rate of correct identifications at the species level for Gram negatives was higher for the differential centrifugation method due to the increased number of isolates from the *Enterobacteriaceae* (87.0%, 60/69) correctly identified by this method versus 84.1% with the rapidBAC pro^®^ II kit (58/69). However, the number of *Pseudomonas aeruginosa* correctly identified at the species level was higher with the commercial kit (100% -4/4- vs. 50.0% -2/4-). Conversely, successful species-level identification was obtained for several Gram-positives using the rapidBAC pro^®^ II kit, especially for *Staphylococcus aureus* (91.7% -11/12- vs. 25.0% -3/12-) or enterococci (91.7% -11/12- vs. 75.0% -9/12-), which added to the global superiority of the commercial kit for the identification of Gram positives at the species level (58.0% -51/88- vs. 44.3% -39/88-) despite its lower rates of correct identifications at this level for streptococci (36.7% -11/30- vs. 46.7% -14/30-). Finally, the percentage of yeast correctly identified by the rapidBAC pro^®^ II kit was also lower (12.5% -1/8- vs. 25.0% -2/8-) but the differential centrifugation method yielded a higher rate of unreliable identified isolates (75.0% -6/8- vs. 62.5% -5/8-).

**Table 4.** Comparison between the performances of the rapidBACpro^®^ test and the in-house Differential Centrifugation on a group of monomicrobial BCs (n=177).

|  |  | n | rapidBACpro II test |  |  |  |  |  | Differential centrifugation method |  |  |  |  |  |
| --- | --- | --- | --- | --- | --- | --- | --- | --- | --- | --- | --- | --- | --- | --- |
|  |  |  | S<br>>2.0 | (%) | 1.7 < S<br>< 1.99 | (%) | S <<br>1.7 | (%) | S<br>>2.0 | (%) | 1.7 < S <<br>1.99 | (%) | S <<br>1.7 | (%) |
| (A) |  |  |  |  |  |  |  |  |  |  |  |  |  |  |
| Overall |  | 177 | 118 | 66.7 | 43 | 24.3 | 16 | 9.0 | 109 | 61.6 | 36 | 20.3 | 32 | 18.1 |
| Gram-Negative (GN) |  | 81 | 66 | 81.5 | 13 | 16.0 | 2 | 2.5 | 68 | 83.9 | 11 | 13.6 | 2 | 2.5 |
| Gram-Positive (GP) |  | 88 | 51 | 58.0 | 28 | 31.8 | 9 | 10.2 | 39 | 44.3 | 25 | 28.4 | 24 | 27.3 |
| (B) |  |  |  |  |  |  |  |  |  |  |  |  |  |  |
| Enterobacterales |  | 69 | 58 | 84.1 | 10 | 14.5 | 1 | 1.4 | 60 | 87.0 | 8 | 11.6 | 1 | 1.4 |
| *Escherichia coli* |  | 35 | 28 | 80.0 | 6 | 17.1 | 1 | 2.9 | 31 | 88.6 | 4 | 11.4 | 0 | 0 |
| *Klebsiella* spp |  | 15 | 14 | 93.3 | 1 | 6.7 | 0 | 0 | 14 | 93.3 | 1 | 6.7 | 0 | 0 |
| *Pseudomonas* |  | 4 | 4 | 100 | 0 | 0 | 0 | 0 | 2 | 50.0 | 2 | 50.0 | 0 | 0 |
| Other GN |  | 8 | 4 | 50.0 | 3 | 37.5 | 1 | 12.5 | 6 | 75.0 | 1 | 12.5 | 1 | 12.5 |
| *S. aureus* |  | 12 | 11 | 91.7 | 1 | 8.3 | 0 | 0 | 3 | 25.0 | 2 | 16.7 | 7 | 58.3 |
| CoN staphylococci |  | 27 | 16 | 59.3 | 11 | 40.7 | 0 | 0 | 12 | 44.5 | 9 | 33.3 | 6 | 22.2 |
| Streptococci |  | 30 | 11 | 36.7 | 11 | 36.7 | 8 | 26.6 | 14 | 46.7 | 9 | 30.0 | 7 | 23.3 |
| Enterococci |  | 12 | 11 | 91.7 | 1 | 8.3 | 0 | 0 | 9 | 75.0 | 2 | 16.7 | 1 | 8.3 |
| Other GP |  | 7 | 2 | 28.6 | 4 | 57.1 | 1 | 14.3 | 1 | 14.3 | 3 | 42.8 | 3 | 42.8 |
| Yeast |  | 8 | 1 | 12.5 | 2 | 25.0 | 5 | 62.5 | 2 | 25.0 | 0 | 0 | 6 | 75.0 |

Of the polymicrobial BCs, 23 cases were included in this head-to-head comparison. The rapidBAC pro^®^ II kit allowed the identification of at least one microorganism in 22/23 cases with score values between 1.75 and 2.55 while the differential centrifugation method provided the identification of one microorganism in 20/23 cases with score values ranging between 1.45 and 2.45. Moreover, the commercial kit allowed the identification of both pathogens in two cases (*S. epidermidis* plus *S. capitis* and *E. faecalis* plus *S. epidermidis*) and the in-house method in one case (*Enterococcus faecium* and *Aeromonas hydrophila* in a BC where *K. pneumoniae* was also present).

In summary, the implementation of the rapidBAC pro^®^ II kit reduced the number of unreliable identifications of microorganisms present in BCs (5.6%; 2.0-9.0% inter-laboratory range) using MALDI-TOF MS. The rate of Gram negatives correctly identified at the species-level was 92.3% and this figure was even higher for certain commonly encounter species (Table 1). Species-level identification of Gram positives directly from BCs, one of MALDI-TOF MS current limitations, was achieved in 72.4% of the cases. Remarkably, 97.3 and 94.7% of the enterococci and *S. aureus* isolates, respectively, were successfully identified using the rapidBAC pro^®^ II kit. However, there is still room for improvement in the case of streptococci (66.3% successful identifications at the species level) or coagulase negative staphylococci (59.5%). The same is true for the other two groups of microorganisms that represent the limitations of MALDI-TOF MS: yeasts, with only 42.9% correct identifications at the species level and polymicrobial BCs since only in 2/30 cases all the species present could be identified.

## DISCUSSION

With the advent of MALDI-TOF MS for rapid identification of microorganisms, several pre-processing methods have been developed for direct identification of pathogens present in BCs. Currently there are three available commercial kits: (i) the Sepsityper^®^ kit –(Bruker Daltonics, Bremen, Germany) (12) (ii) the Vitek^®^ MS Blood Culture Kit (bioMérieux, Inc., Marcy l’Etoile, France) (13) and (iii) the rapidBAC pro^®^ II kit (Nittobo Medical Co., Tokyo, Japan) (14). In addition, several in-house methods have been developed for this purpose (2, 6-7, 15). The Sepsityper^®^ kit has been extensively evaluated and successful, accurate identifications at the species level over 80.0% have been reported (12). However, evaluations of the rapidBAC pro^®^ II kit performance on BCs are still scarce. Tsuchida et al. (14) have analyzed a collection of 193 monomicrobial BCs and demonstrated an overall correct microbial identification of 80.8% and 99.5% with score values cut-offs at ≥2.0 and ≥1.7, respectively. The same study observed that all Gram positives were correctly identified at the species level with score values ≥1.7 and in 69.4% of the cases with score values ≥2.0. In a more recent study including 199 BCs, Kayin et al. showed that the rapidBAC pro^®^ II kit yielded overall 87.4% correct species-level identification (score value ≥1.7) using the standard module of the biotyper system (Bruker Daltonics). This rate increased to 94.4% when the specific Sepsityper module for the Sepstityper kit (score value ≥1.6) was applied. Gram positives were accurately identified at the species level in 91.5% of the cases using the Sepsityper module (16).

In our study, 761 monomicrobial BCs were analyzed in four laboratories using the rapidBAC pro^®^ II kit. We identified 80.0% of the isolates correctly at the species level with score values ≥2.0 and 94.4% correctly at the genus level with score values ≥1.7. These results are in agreement with those obtained by Tsuchida *et al*. (2018) and Kayin *et al*. (2019). The enterococci (97.3%) and *S. aureus* (94.7%) were correctly identified with scores ≥2.0 more often than other Gram positives such as streptococci (66.3%) and coagulase negative staphylococci (59.5%). Regarding the Gram negatives, 92.3% of the isolates were correctly identified to species level with score values ≥2.0. Finally, 42.9% of yeasts included in the study were accurately identified with score values ≥2.0. The low capacity of MALDI-TOF MS to identify yeasts from blood cultures at species level was already reported (17), likely due to the presence of a specific cell wall and the interference of blood components in the protein spectra (18). In this study, 38.1% of the yeasts were unreliably identified even when applying score values <1.7 as cut-off, showing one of the limitations of the rapidBAC pro^®^ II kit.

A subset of 560 BCs was analyzed with the rapidBAC pro^®^ II and the Sepsityper^®^ kit in a head-to-head comparison. High rates of Gram negatives were correctly identified by both kits: 90.5% with the rapidBAC pro^®^ II kit and 85.8% with the Sepsityper^®^ kit (score values ≥2.0). The rate of correct species assigned was higher than 90% for *E. coli, K. pneumoniae* and other *Enterobacteriaceae* using both kits, as previously observed (12). However, 100% of the *P. aeruginosa* isolates were identified with score values ≥2.0 using the rapidBAC pro^®^ II kit vs. 70.0% with the Sepsityper^®^ kit. Higher differences were observed for Gram positive bacteria: 68.7% with the rapidBAC pro^®^ II kit (score values ≥2.0) and 59.7% with the Sepsityper^®^ kit. Within this group, major differences were observed in the identification of coagulase-negative staphylococci (55.9% with the rapidBAC pro^®^ II kit vs. 43.2% using the Sepsityper^®^ kit -score values ≥2.0-) and enterococci (96.7% with the rapidBAC pro^®^ II kit vs. 76.7% using the Sepsityper^®^ kit -score values ≥2.0-). Errors in the identification of coagulase negative bacteria present in blood culture pellets have been reported (7). However, no misidentifications with any of the applied methods have been detected in this study.

The performance of two in-house pre-processing methods were also compared with rapidBAC pro^®^ II kit. The method containing a lysis step with ammonium chloride showed lower accuracy than the rapidBAC pro^®^ II kit when applied to the different microbial pellets: overall species-level identifications -89.0% vs 39.8%-, Gram negatives -96.7% vs 53.8%-, Gram positives -83.6% vs 26.9%- and yeasts 66.7% vs 50.0%-. These results are lower than those previously obtained by the same research group (83.0% and 42.0% of correct species level identifications for Gram negative and Gram positive bacteria, respectively) (7). Differences may be due to the different set of bacterial species included in this study. Regarding the differential centrifugation method, the overall rate and the amount of correct identifications of Gram positives was higher for the rapidBAC pro^®^ II kit (58.0% vs. 44.3%, respectively) but Gram negatives were identified in a more accurate way by the in-house method (83.9% vs. 81.5%).

Among 27 polymicrobial BCs, we could identify both pathogens twice (7.4%) with the rapidBAC pro^®^ II kit and once (3.7%) with Sepsityper^®^ kit. These rates are lower than those reported by Scohy *et al*. (2018), who identified two pathogens in 36.8% of the cases by applying the specific Sepsityper module from the Compass IVD software -Bruker Daltonics- (19). Further evaluation of the rapidBAC pro^®^ II kit performance coupled to the Sepsityper module will be carried out in order to elucidate if the combination of the improved sample preparation method and the *ad-hoc* method for spectra acquirement may improve further the identification of polymicrobial BCs.

In summary, the evaluated commercial kit allowed a high rate of correct identifications of Gram negative and Gram positive bacteria. They were less effective for the identification of yeasts. Results obtained were consistent in the four laboratories where the kit was tested, showing a reduced number of not reliable identification and a high rate of Gram positive bacteria reliably identified at the species level. Hands-on time was calculated as 30 to 40 minutes for 1 to 10 BCs, which enables the implementation of this methodology in the laboratory routine in a standardized way. Moreover, the kit is manufactured independently from the mass spectrometry systems, which provides flexibility to the MALDI-TOF MS user regarding the optimized use of available reagents. In addition, we confirmed the robustness of the MALDI-TOF MS approach, since we observed no discordant results even when applying different pellet preparation methods.

## CONFLICT OF INTERESTS

Nittobo Medical (Tokyo, Japan) has provided the reagents for this study. The company had no role in the study design, data collection and analysis, decision to publish, or preparation of the manuscript.

## ACKNOWLEDGEMENTS

BRS is supported by the Miguel Servet Program (MSII19/00002) and MO by the Joan Rodés Program (JR18/00006). Both contracts are funded by the Health Research Fund (FIS) of the Carlos III Health Institute (ISCIII-MICINN), Spain, partially financed by the by the European Regional Development Fund (FEDER) ‘A way of making Europe’. The funders The funders played no role in the study design, data collection and analysis, decision to publish or preparation of the manuscript.

## REFERENCES

1. Patel R. MALDI-TOF MS for the diagnosis of infectious diseases. 2015. Clin Chem 61:100–11. doi: 10.1373/clinchem.2014.221770.

2. Rodríguez-Sánchez B, Cercenado E, Coste AT, Greub G. Review of the impact of MALDI-TOF MS in public health and hospital hygiene, 2018. 2019. Euro Surveill 24:1800193. doi: 10.2807/1560-7917.ES.2019.24.4.1800193.

3. Opota O, Croxatto A, Prod’hom G, Greub G. 2015. Blood culture-based diagnosis of bacteraemia: state of the art. Clin Microbiol Infect 21:313–22. doi: 10.1016/j.cmi.2015.01.003

4. Oviaño M, Bou G. 2018. Matrix-Assisted Laser Desorption Ionization-Time of Flight Mass Spectrometry for the Rapid Detection of Antimicrobial Resistance Mechanisms and Beyond. Clin Microbiol Rev 32:e00037–18. doi: 10.1128/CMR.00037-18.

5. Leibovici L, Shraga I, Drucker M, Konigsberger H, Samra Z, Pitlik SD. 1998. The benefit of appropriate empirical antibiotic treatment in patients with bloodstream infection. J Intern Med 244: 379–386.

6. March-Rosselló GA, Muñ oz-Moreno MF, García-Loygorri-Jordán de Urriés MC, Bratos-Pérez MA. 2013. A differential centrifugation protocol and validation criterion for enhancing mass spectrometry (MALDI-TOF) results in microbial identification using blood culture growth bottles. Eur J Clin Microbiol Infect Dis 32:699–704. doi: 10.1007/s10096-012-1797-1

7. Prod’hom G, Bizzini A, Durussel C, Bille J, Greub G. 2010. Matrix-assisted laser desorption ionization-time of flight mass spectrometry for direct bacterial identification from positive blood culture pellets. J Clin Microbiol 48:1481–3. doi: 10.1128/JCM.01780-09.

8. Bidart M, Bonnet I, Hennebique A, Kherraf ZE, Pelloux H, Berger F, et al. 2015. An in-house assay is superior to Sepsityper for direct matrix-assisted laser desorption ionization-time of flight (MALDI-TOF) mass spectrometry identification of yeast species in blood cultures. J Clin Microbiol 53:1761–4. doi: 10.1128/JCM.03600-14.

9. Yonetani S, Ohnishi H, Ohkusu K, Matsumoto T, Watanabe T. 2016. Direct identification of microorganisms from positive blood cultures by MALDI-TOF MS using an in-house saponin method. Int J Infect Dis 52:37–42. doi: 10.1016/j.ijid.2016.09.014.

10. Idelevich EA, Schüle I, Grünastel B, Wüllenweber J, Peters G, Becker K. 2014. Rapid identification of microorganisms from positive blood cultures by MALDI-TOF mass spectrometry subsequent to very short-term incubation on solid medium. Clin Microbiol Infect 20(10):1001–6. doi: 10.1111/1469-0691.12640.

11. Florio W, Cappellini S, Giordano C, Vecchione A, Ghelardi E, Lupetti A. 2019. A new culture-based method for rapid identification of microorganisms in polymicrobial blood cultures by MALDI-TOF MS. BMC Microbiol 19(1):267.

12. Schieffer KM, Tan KE, Stamper PD, Somogyi A, Andrea SB, Wakefield T, et al. 2014. Multicenter evaluation of the Sepsityper™ extraction kit and MALDI-TOF MS for direct identification of positive blood culture isolates using the BD BACTEC™ FX and VersaTREK^®^ diagnostic blood culture systems. J Appl Microbiol 116:934–41.

13. Broyer P, Perrot N, Rostaing H, Blaze J, Pinston F, Gervasi G, et al. 2018. An Automated Sample Preparation Instrument to Accelerate Positive Blood Cultures Microbial Identification by MALDI-TOF Mass Spectrometry (Vitek®MS). Front Microbiol 9:911. doi: 10.3389/fmicb.2018.00911.

14. Tsuchida S, Murata S, Miyabe A, Satoh M, Takiwaki M, Ashizawa K, et al. 2018. Application of the biocopolymer preparation system, rapid BACpro® II kit, for mass-spectrometry-based bacterial identification from positive blood culture bottles by the MALDI Biotyper system. J Microbiol Methods 152:86–91.

15. Florio W, Morici P, Ghelardi E, Barnini S, Lupetti A. 2018. Recent advances in the microbiological diagnosis of bloodstream infections. Crit. Rev. Microbiol 44(3):351–370.

16. Kayin M, Mert B, Aydemir S, Özenci V. 2019. Comparison of rapid BACpro® II, Sepsityper® kit and in-house preparation methods for direct identification of bacteria from blood cultures by MALDI-TOF MS with and without Sepsityper® module analysis. Eur J Clin Microbiol Infect Dis 38:2133–2143.

17. Azrad M, Keness Y, Nitzan O, Pastukh N, Tkhawkho L, Freidus V, et al. 2019. Cheap and rapid in-house method for direct identification of positive blood cultures by MALDI-TOF MS technology. BMC Infect Dis 19:72.

18. Buchan BW, Ledeboer NA. 2013. Advances in Identification of Clinical Yeast Isolates by Use of Matrix-Assisted Laser Desorption Ionization–Time of Flight Mass Spectrometry. J Clin Microbiol 51:1359–1366.

19. Scohy A, Noël A, Boeras A, Brassinne L, Laurent T, Rodriguez-Villalobos H, et al. 2018. Evaluation of the Bruker® MBT Sepsityper IVD module for the identification of polymicrobial blood cultures with MALDI-TOF MS. Eur J Clin Microbiol Infect Dis 37:2145–2152.

